## Supplementary figures and images for "*In silico* repurposed drugs against monkeypox virus"

### Supplementary figure 1

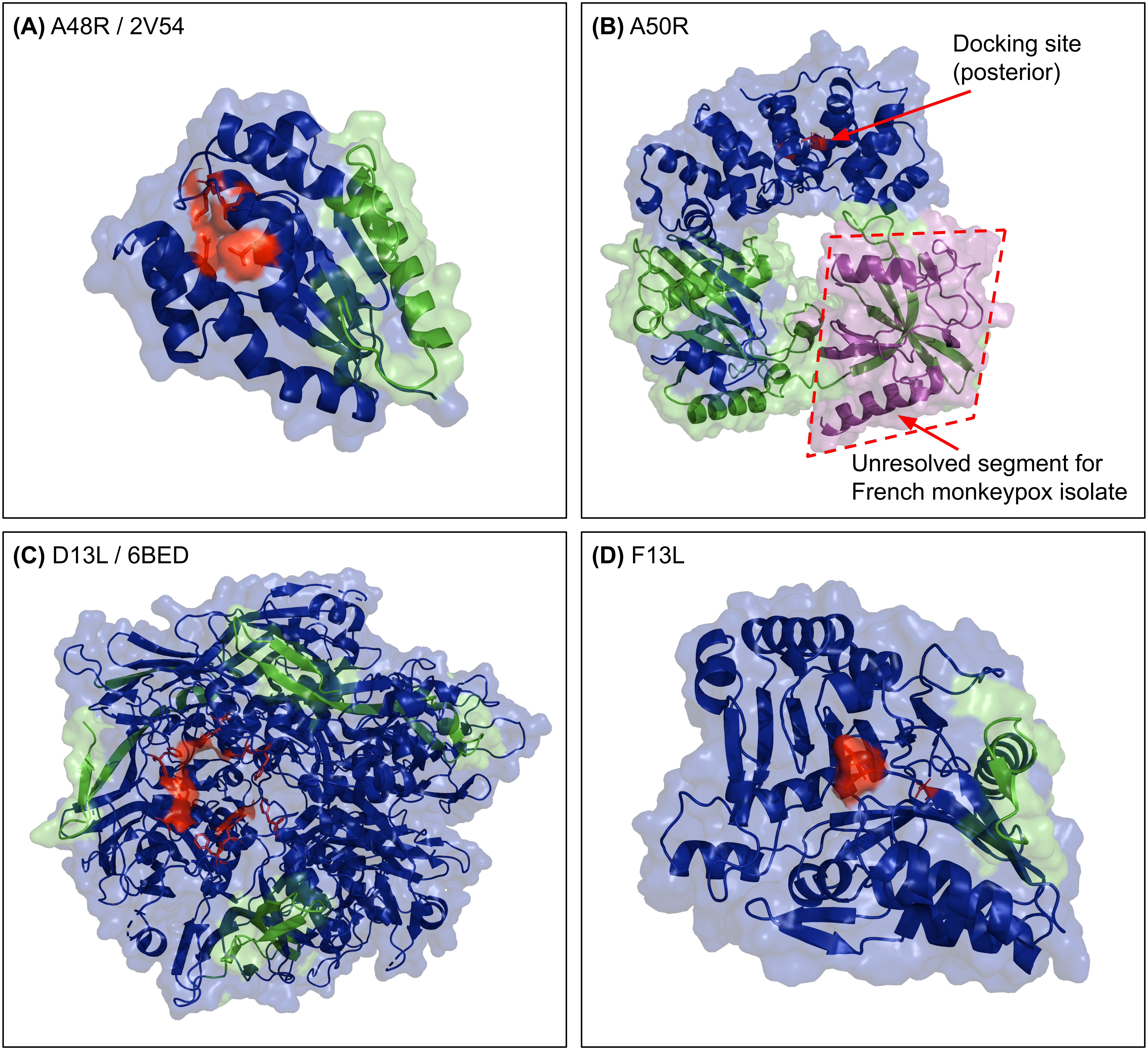

### Supplementary figure 2

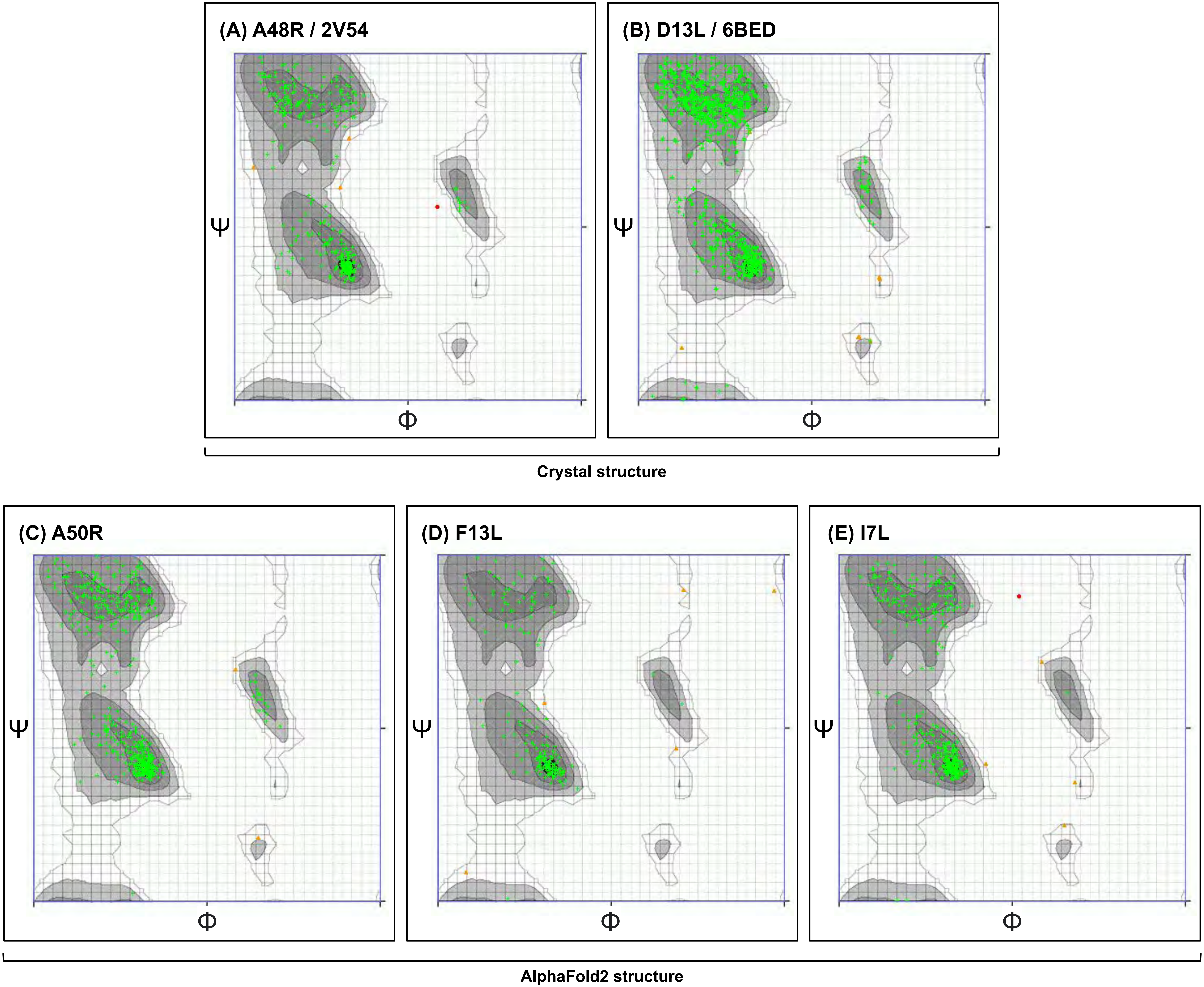

### Supplementary figure 3

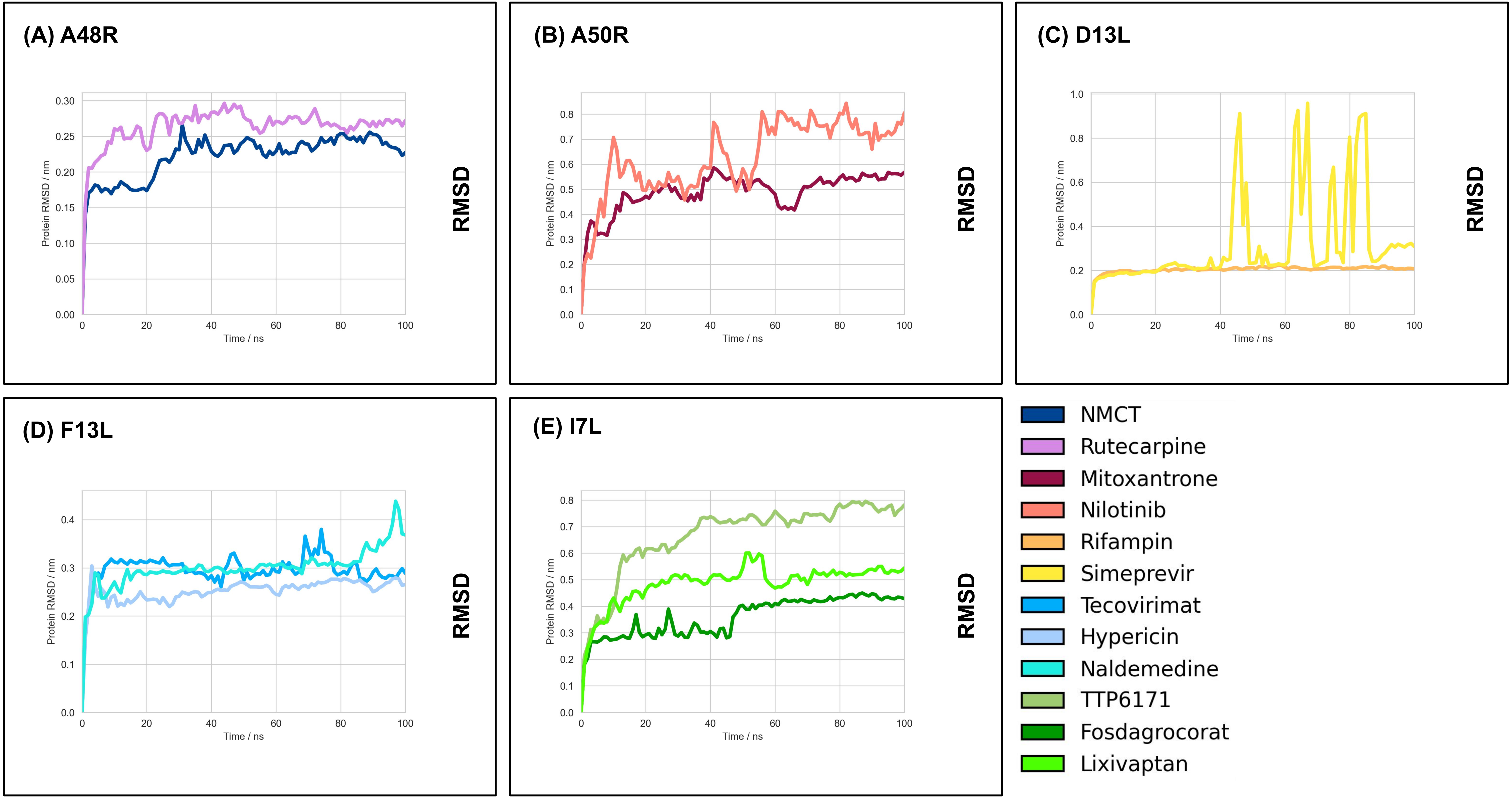

### Supplementary figure 4

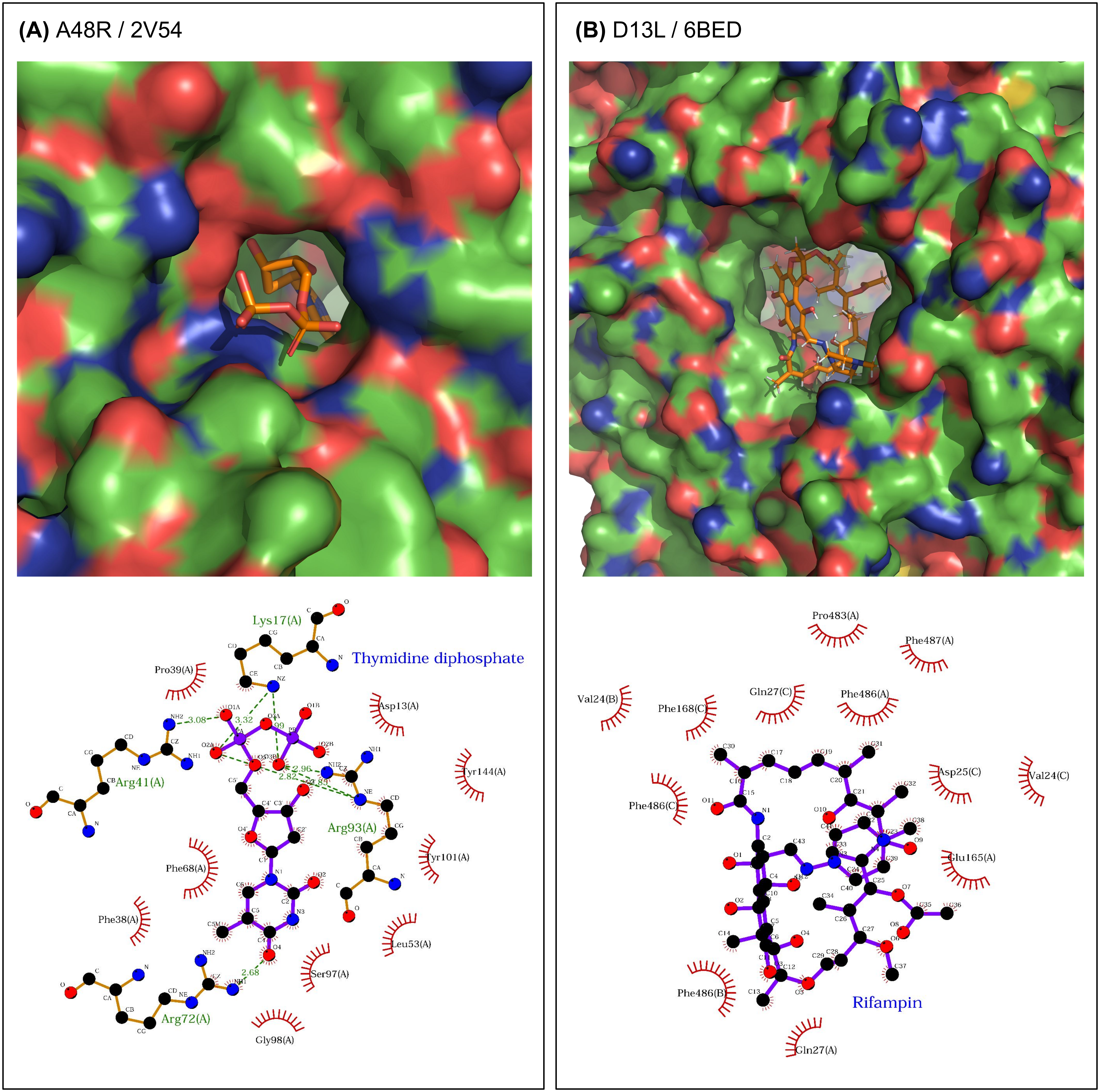

### Supplementary figure 5

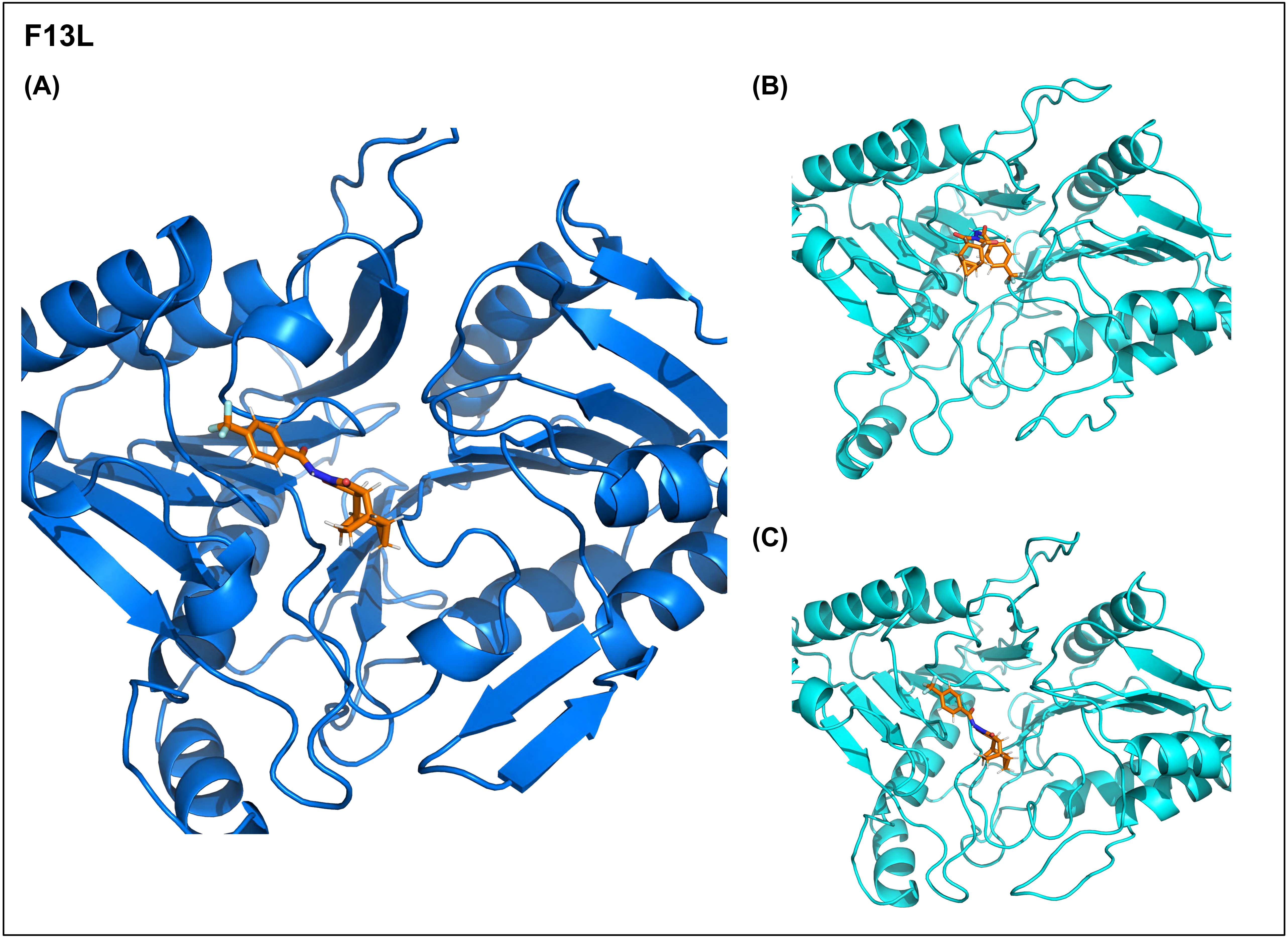
